# Large-language-model-based antibiotic resistance gene prediction and resistomes mining in cyanobacterial blooms

**DOI:** 10.1101/2025.09.03.674132

**Authors:** Haimeng Li, Miao Qi, Xin Wang, Honglei Liu, Qiuying Yang, Hui Chen, Rui beijingJiang, Ting Chen, Yuqing Yang, Congmin Zhu

**Affiliations:** State Key Laboratory of Networking and Switching Technology, Beijing University of Posts and Telecommunications, Beijing, China; School of Biomedical Engineering, Capital Medical University; Beijing, China; Department of Ultrasound, Peking Union Medical College Hospital, Beijing, China; Chinese Academy of Medical Sciences, Peking Union Medical College, Beijing, China; Beijing Key Laboratory of Fundamental Research on Biomechanics in Clinical Application, Capital Medical University, Beijing, China; Bioinformatics Division and Center for Synthetic & Systems Biology, Beijing National Research Center for Information Science and Technology; Department of Automation, Tsinghua University, Beijing, China; Institute for Artificial Intelligence and Department of Computer Science and Technology, Tsinghua University, Beijing, China

**Keywords:** cyanobacterial aggregate, metagenomics, antibiotic resistance genes, large language model

## Abstract

The rapid spread of antibiotic resistance genes (ARGs) in aquatic ecosystems poses a serious public health threat. Conventional ARG detection methods, including sequence-alignment and machine learning, are limited by high false-negative or false-positive rates, especially due to sequence diversity and class imbalance in metagenomic data. Moreover, few tools offer comprehensive solutions for ARG identification, classification, and functional annotation. To address these limitations, we developed ESMARG, a novel protein language model framework based on ESM1v, for accurate ARG identification, classification, and annotation. Trained with both ARG and abundant non-ARG sequences from real-world metagenomes to enhance robustness and reduce false positives, ESMARG significantly outperformed traditional alignment-based (BLAST, DIAMOND) and recent deep learning models (ARGNet, ARG-SHINE), achieving 0.998 precision, 0.939 recall, and 0.968 F1-score for ARG identification. Functional and mechanism classification modules also demonstrated high accuracy (0.986 and 0.987) and computational efficiency. We applied ESMARG to analyze 26 cyanobacterial aggregate (CA) metagenomes collected from Lake Taihu across a full annual cyanobacterial harmful algal bloom (CyanoHAB) cycle. A total of 110 ARGs, spanning 24 drug classes and multiple resistance mechanisms, were detected, with seasonal shifts in ARG abundance and composition. Strong correlations were identified between CA resistomes and microbial community structure, especially with cyanobacteria. Further, ARG abundance was shaped by both biotic factors—such as cyanobacterial dominance and bacterial functional profiles—and abiotic variables, including biochemical oxygen demand and water temperature. Our findings demonstrate the power of ESMARG for high-resolution ARG profiling and reveal complex ecological interactions between resistomes, microbiomes, and environmental factors in CyanoHAB.

**Highlights:**

1. Developed ESMARG model outperforms BLAST/other deep learning models in ARG detection
2. ARG showed seasonal dynamics in cyanobacterial metagenomes across algal bloom cycle
3. ARG was strongly correlate with microbiome structure and cyanobacteria dominance
4. ARG abundance was shaped by cyanobacteria, bacterial functions, BOD and temperature

## 1. Introduction

Antibiotic resistance - the ability of bacteria to survive and replicate in the presence of an antibiotic [1]- is an escalating global public health threat [2]. Many pathogens have acquired resistance to major antibiotics, with some evolving multidrug resistance, making infections increasingly difficult or even impossible to treat [3, 4]. The widespread use of antibiotics in human medicine, agriculture, and animal husbandry has created environmental reservoirs for antibiotic-resistant bacteria (ARB) and their associated antibiotic resistance genes (ARGs), especially in aquatic ecosystems [5, 6]. These ARGs disseminate through surface water, sewage, drinking water, and natural water bodies [7], where they can be horizontally transferred to pathogens, reducing the efficacy of antibiotic treatments and posing significant risks to human and ecosystem health [8]. Increasing evidence also suggests that microbial community structure plays a key role in shaping ARG profiles in freshwater systems [9, 10].

Accurate identification of ARGs is essential for monitoring and mitigating antibiotic resistance. Current ARG detection methods generally fall into two main categories: sequence-alignment-based and machine-learning-based approaches. Traditional tools like BLAST [11], DIAMOND [12], and Bowtie2 [13] match sequences to reference databases (e.g., CARD [14], SARG [15]) and are widely used to identify known ARGs. However, these tools heavily depend on the quality and integrity of the reference databases, which may limit their ability to recognize newly emerging or rare ARGs.

Machine learning tools like HMMER [16] uses hidden Markov models to detect remote homology, while TGC-ARG [17] incorporates secondary structure prediction. Deep learning methods have further advanced the field. DeepARG [18], uses MLP to classify ARGs based on sequence length, while HMD-ARG [19] and ARG-SHINE [20] employ a CNN architecture for ARGs identification. ARGNet [21] uses attention-based models for identifying both known and novel ARGs without relying on sequence alignment. Despite advances, current models struggle with class imbalance due to the overwhelming abundance of non-ARG sequences in metagenomes, leading to high false positives and limited utility. Furthermore, few tools offer an integrated pipeline for comprehensive ARG detection, classification (by drug and mechanism), and annotation. To overcome these limitations, a new LLM-based framework for ARG identification and classification was developed in our study.

Cyanobacterial Harmful Algal Blooms (CyanoHABs) represent a growing threat to freshwater ecosystems worldwide, with eutrophication being a primary driver [22, 23]. In lakes like Lake Taihu, excessive nutrient loadings have caused CyanoHABs to become frequent and intense over the past few decades [24, 25]. These blooms not only degrade water quality but also disrupt local aquatic ecosystems and pose risks to human health through contamination of drinking water sources. Cyanobacterial Aggregates (CA), the basic unit wherein cyanobacteria and bacteria coexist in close contact, is the primary form of cyanobacteria during blooms. Cyanobacteria, as autotrophic organisms, provide food to their attached bacterial communities by secreting extracellular organic matter, while the bacteria supply essential nutrients such as nitrogen, phosphorus, sulfur, and trace elements to the cyanobacteria [26, 27]. Since the mutualistic relationships were reported to be exist between cyanobacteria and bacteria in CAs at both taxonomic and gene levels [28, 29], ARG composition of bacteria is likely to strongly be shaped in response to CyanoHABs, or they can influence the outbreak of CyanoHABs. However, to the best of our knowledge, the role of bacterial resistomes (i.e., collections of ARGs) [30] during cyanobacterial blooms is remained unknown. Strong inter actions between the ARGs, microbial assemblages and the cyanobacterial blooms need to be understood in the broader context of managing ARGs in aquatic ecosystems.

To address these challenges, we present ESMARG, a large language model (LLM)-based framework built on protein language models (ESM1v)[31], capable of accurate ARG detection, functional classification, mechanism prediction, and annotation (Fig.1). To reduce the false positive rate, plenty of non-ARG sequences from real metagenomic sequencing samples have been included to refine the model. Then, we apply ESMARG to analyze the distribution of ARGs in CA metagenomes from Lake Taihu covering a full CyanoHAB cycle, aiming to (1) monitor the variations of ARG composition throughout different cyanobacterial bloom periods; (2) explore the relationships between resistomes and microbiome in CAs; and (3) reveal the contribution of biotic and abiotic factors to the variations of ARGs in CAs.

**Figure 1.**
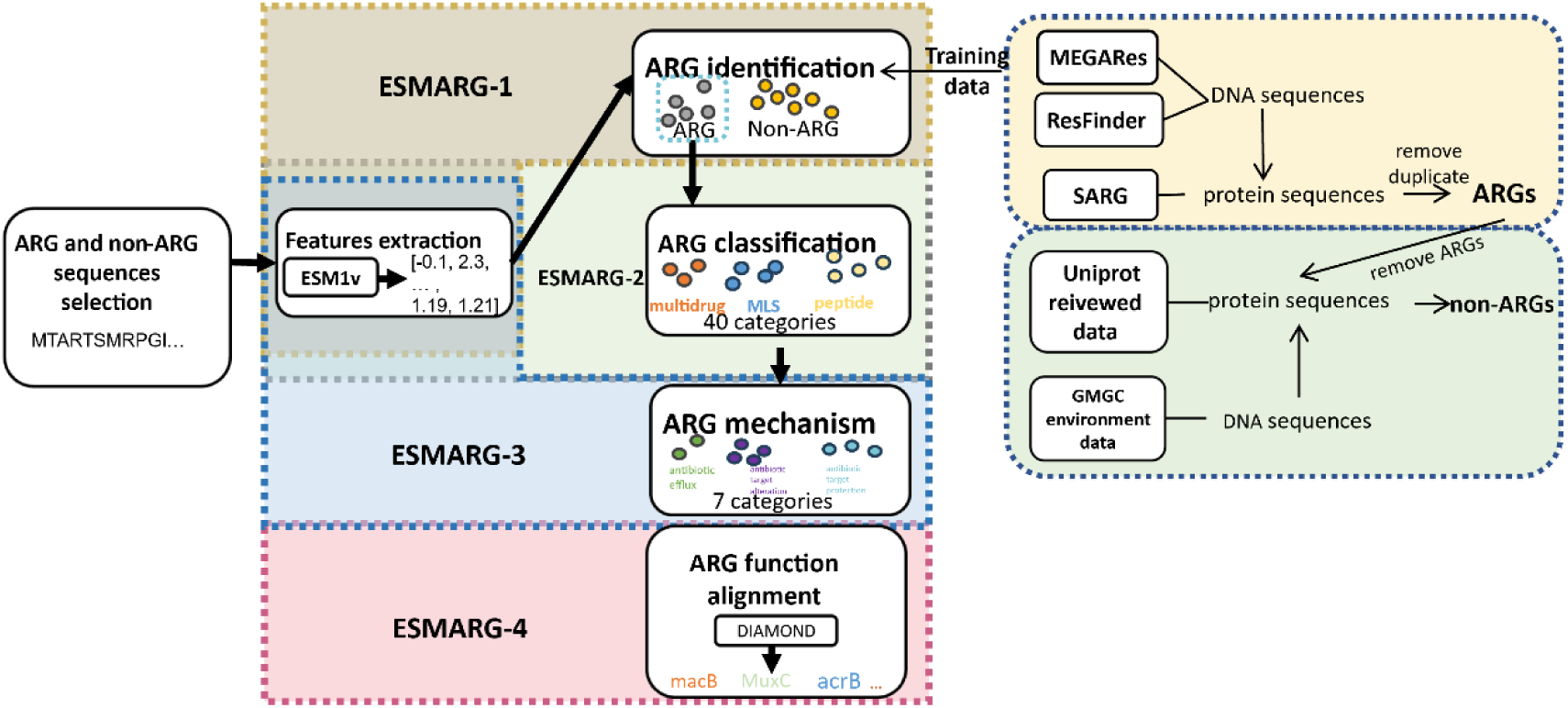
Schema of large-language-model-based antibiotic resistance gene prediction method. The ESMARG-1 model is designed to classify whether sequences are ARG sequences or no and its subsequent model ESMARG-2 is for classifying the categories of ARGs, ESMARG-3 is for classifying antibiotic mechanisms of ARGs and ESMARG-4 is for aligning the functions of ARGs using DIAMOND. ARG sequences were from the databases: MEGARes, ResFinder, and SARG. Negative samples are selected from UniProt reviewed data and GMGC environmental data.

## 2. Methods

### 2.1 Data processing with self-constructed ARG dataset

A novel model, ESMARG, consisted of four modules: ESMARG-1, ESMARG-2, ESMARG-3, and ESMARG-4 was proposed. ESMARG-1 was designed to classify sequences as ARGs or non-ARGs. ESMARG-2 categorizes ARGs into functional groups, while ESMARG-3 identifies their resistance mechanisms. ESMARG-4 performs detailed annotation of ARGs via sequence alignment. The positive samples, 15624 ARG sequences, were extracted from several publicly available databases: CARD [14], SARG [15], MEGARes [32] and ResFinder [33]. Negative samples for ESMARG-1 were selected from two primary datasets: the UniProt database [34] and GMGC [35], which included the real environmental metagenomic sequences of food, livestock, compost, soil, and water. DIAMOND was employed to remove known ARGs from GMGC sequences using different e-value thresholds (e.g. 1e-5). Detailed descriptions of the data processing procedures are provided in the appendix.

### 2.2 ESMARG Model architecture

To capture the semantic information of ARG sequences for accurate identification, the protein LLM, ESM1v[31], was employed as the foundation and a comprehensive pipeline named ESMARG was developed, which was capable of identifying potential ARGs in ORF sequences, classifying their functions and their mechanisms, and annotating the sequences. Building on the ESM1v backbone, a multi-layer perceptron (MLP) was utilized to fine-tune the model. Detailed hyperparameter settings of the model are provided in the appendix.

### 2.3 Evaluation and comparison based on external Oxford ARG datasets

The extra test dataset developed by Oxford database [36] was selected to evaluate the performance of ESMARG-1, ARGNet and BLAST. ARGs within Oxford database had no sequence annotation information, so the first step was to download the sequence corresponding to each ARG using UniProt’s API. A total of 420 ARG sequences were selected. For the ESMARG-1, to identify whether a new sequence was an ARG the combination of negative samples was used:30,000 from UniProt and 40,000 from GMGC.

### 2.4 Performance evaluation of four modules from ESMARG

The performance of ESMARG-1 was evaluated with three primary metrics, precision, recall, and F1-score. Other ARG identification methods such as ARGNet, BLAST and DIAMOND were chosen as the baseline. Besides, ROC curves were drawn to show overall predictive performance of different methods.

In ESMARG-1, the training process was conducted multi times with different negative samples (70,000,100,000, and 200,000) from the UniProt database to evaluate the influence of non-ARGs on model efficiency. Besides, extra negative samples from the GMGC were also included for training, enhancing the robustness of the module to false positive problem. For comparison, results of BLAST and DIAMOND with an e-value threshold of 1e-5, identity of 50% and the max target sequence of 1 were analyzed, deep learning model ARGNet[21] was also evaluated with the same test datasets.

For ESMARG-2 and ESMARG-3, accuracy and weighted-F1 score were used to evaluate the performance of ARG categories classification and ARG resistance mechanism classification. Weighted-F1 score defined as,

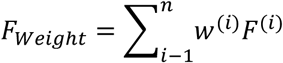

which considered the issue of sample imbalance. To compare with other models regarding the classification of the categories of ARGs, BLAST, DIAMOND were selected to calculate accuracies and F1 including an e-value threshold of 1e-5 and a sequence identity threshold of 50%, deep learning model ARG-SHINE [20] was also evaluated with the same test datasets. ESMARG-3 compared weighted F1 with DIAMOND and BLAST (the same value in ESMARG-2 comparison).

### 2.5 ARG analysis in CA samples

For the ARG analysis of CA samples, we utilized the ESMARG-1 module, which was trained on 30,000 negative samples from UniProt and 40,000 negative environmental samples from GMGC. The model achieved high performance metrics, with a precision of 0.998, recall of 0.956, and an F1-score of 0.977. Sequences predicted to be potential antibiotic resistance genes (ARGs) were selected based on prediction scores exceeding 0.6.

For these candidate ARG sequences, we employed ESMARG-2 to classify their categories and ESMARG-3 to determine their antibiotic resistance mechanisms. To evaluate the results obtained from ESMARG-1, ESMARG-2, and ESMARG-3, high-scoring protein sequences are selected for further analysis. Sequences classified by ESMARG-1 were aligned with the DeepARG database using ESMARG-4 to investigate their functional annotations.

To evaluate the distribution of ARGs in AS samples, the abundance of ARGs was normalized and represented as “ARG copies per cell” following the published paper [37] using the following equation.:

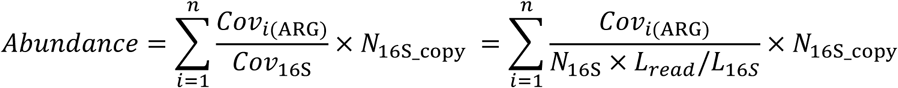

where *n* denotes the number of annotated ARGs, *Cov*_*i*(ARG)_ represents the coverage of a gene identified as ARG, *N*_16S_ corresponds the count of 16S rRNA gene sequences detected within the metagenomic data, *L*_*read*_ indicates the read length (bp), *L*_16*S*_ refers to the mean length of 16S rRNA genes (1432 bp) as provided in Greengenes database, and *N*_16S_copy_ signifies the average 16S rRNA gene copy number per cell in the community. The value of *N*_16S_copy_ was calculated as the abundance-weighted mean of the 16S rRNA gene copy number, wherein the copy number for each genus was estimated using the rrnDB database based on its closest relatives with documented rRNA copy numbers. It should be noted that the normalized ARG abundance (gene copies per cell) is contingent upon the algorithms employed for identifying ARGs and 16S rRNA genes. Given the potential for false positives and false negatives, the resulting ARG abundance may not accurately reflect the true ARG levels within the community. Assuming that the estimates across different samples are subject to a similar degree of bias, we can then compare the ARG abundance among samples and investigate the underlying mechanisms that shape the resistomes.

Detailed statistical analysis methods are provided in the appendix.

## 3. Results

### 3.1 Performance of ARG identification of ESMARG compared to other methods

Performance of ESMARG was evaluated from three aspects: identifying whether a sequence was an ARG, classifying the function and mechanism categories of ARGs. As shown in Fig. 2A, ESMARG-1, one module of ESMARG for ARG identification, achieved higher precision (0.988), recall (0.939), and F1-score (0.962) when lots of non-ARGs from real environmental samples from GMGC database were extracted and included into the training dataset. By increase the ratio of negative samples in training process, the false positive problem of the deep learning model can be improved. ROC curves of ESMARG-1 and other methods were shown in Fig. 2B, and ESMARG-1 (with a classification threshold 0.6, shown in Table 1) demonstrated superior performance in precision (0.998), recall (0.939), and F1-score (0.968) compared to BLAST and DIAMOND. Deep learning model ARGNet (precision 0.778, recall 0.916, F1 0.842) was also compared on the same test set[21]. We further evaluated the proposed method’s performance using the Oxford ARG database with manually collected ARGs [36]. In extra test dataset consisted of 420 ARGs from Oxford database, 302 sequences were predicted as ARGs by ESMARG-1, 268 sequences by BLAST and 296 sequences by ARGNet, which verify the efficiency of ESMARG-1 again.

**Figure 2.**
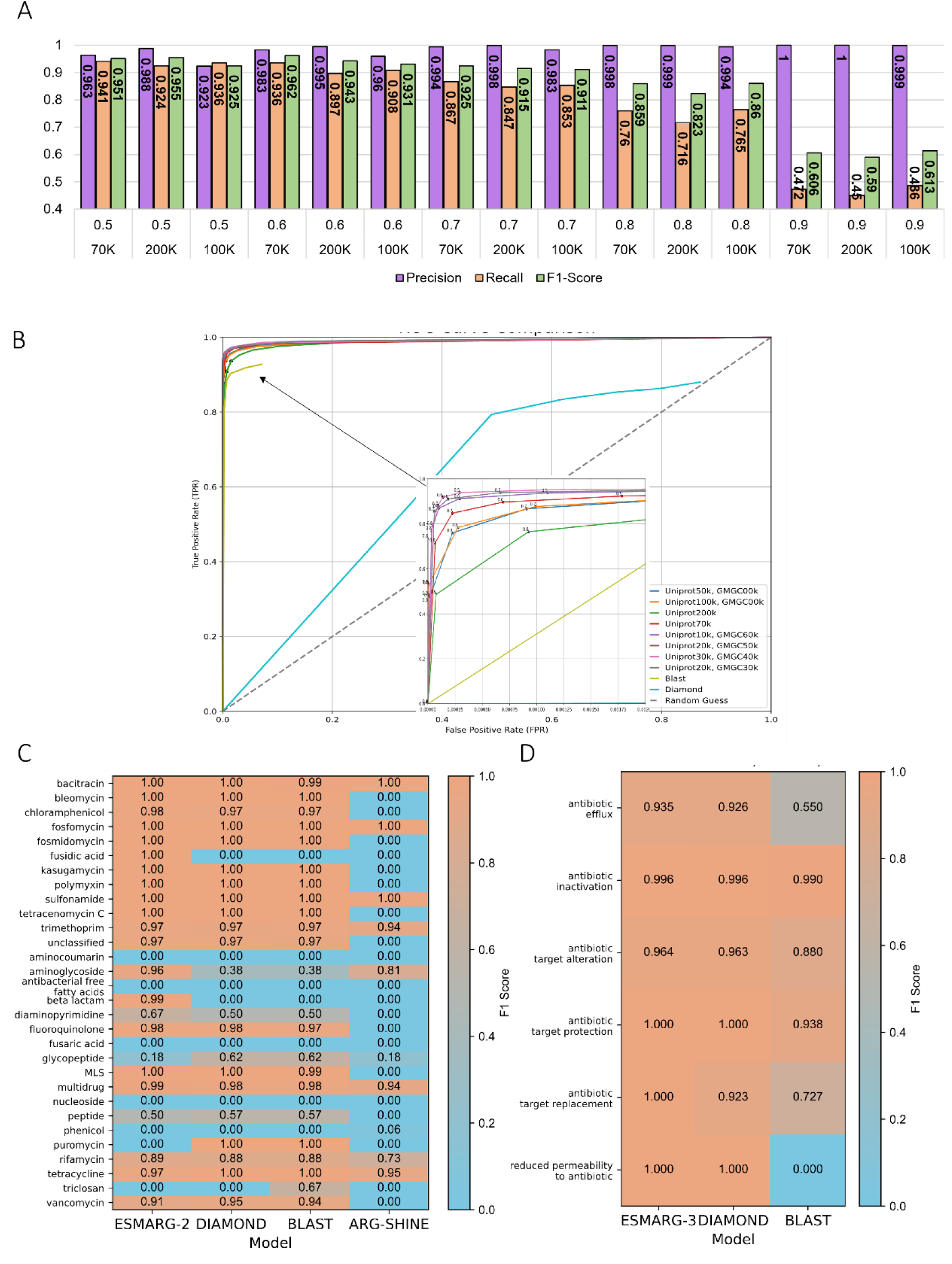
ESMARG’s predictive performance on test dataset. **A)** Precisions, recalls, and F1 values of ESMARG-1 with different thresholds and number of negative samples. (0.5 means a threshold of 0.5 was used to determine predicted sequences as ARGs, and 70k refers to the number of negative samples.) **B)** Comparison of ROC curves for BLAST, DIAMOND, and the ESMARG-1 model with varying negative sample sizes. (Uniprot50k: 50,000 negative samples from Uniprot; GMGC60k: 60,000 negative samples from GMGC.) **C)** F1 scores of ESMARG-2, DIAMOND, BLAST and ARG-SHINE for different categories. Some categories don’t have scores since they have a few samples and don’t have test set. **D)** F1 scores of ESMARG-3, BLAST and DIAMOND were assessed for different antibiotic mechanisms.

**Table 1.** Influence of Negative Sample Selection Strategy on ARG Recognition Performance.

| Negative samples | Precision | Recall | F1-score |
| --- | --- | --- | --- |
| UniProt30k<br>+GMGC40k | 0.998 | 0.939 | 0.968 |
| Uniprot10k<br>GMGC40k | 0.996 | 0.941 | 0.968 |
| Uniprot50k<br>GMGC20k | 0.986 | 0.950 | 0.968 |
| Uniprot20k<br>GMGC30k | 0.996 | 0.939 | 0.967 |
| Uniprot10k<br>GMGC60k | 0.993 | 0.938 | 0.964 |
| Uniprot70k | 0.988 | 0.939 | 0.963 |
| Uniprot50k | 0.948 | 0.915 | 0.948 |
| Uniprot200k | 0.995 | 0.897 | 0.943 |
| Uniprot100k | 0.960 | 0.908 | 0.931 |

Uniprot30k: 30,000 negative samples from Uniprot, GMGC40k: 40,000 negative samples from GMGC.

### 3.2 Predictive validity of function and mechanism classification of ESMARG

The module ESMARG-2 (function prediction of ARGs) and ESMARG-3(ARG resistance mechanism classification) achieving superior results in both accuracy (0.986 and 0.906) and weighted-F1 scores (0.981 and 0.992) compared to BLAST and DIAOMND, while ESMARG-2 also outperformed the state-of-the-art model ARG-SHINE (Table 2). Specifically, high prediction accuracy for ESMARG-2 was observed for categories such as MLS, bacitracin, and fosmidomycin, while lower accuracy was noted for aminoglycoside and nucleoside categories (Fig. 2C). Mechanisms in ESMARG-3 such as reduced permeability to antibiotic target protection, antibiotic target alteration and antibiotic target protection replacement achieved F1-scores exceeding 0.95, whereas the classification performance for antibiotic efflux was comparatively lower (Fig. 2D). ESMARG demonstrated a significant improvement in computational efficiency compared to BLAST and DIAMOND when processing the same set of sequences, representing a 10-fold speedup (Fig. S1D).

**Table 2.** The accuracies and weighted-F1 scores of ESMARG-2, ESMARG-3, DIAMOND, BLAST and ARG-SHINE on the test set.

| Metrics | Model | Function for ARG model |  |
| --- | --- | --- | --- |
|  |  | ARG category | ARG mechanism |
| Accuracy | ESMARG | 0.986 | 0.906 |
|  | DIAMOND | 0.982 | 0.915 |
|  | BLAST | 0.981 | 0.918 |
|  | ARG-SHINE | 0.808 | - |
| Weighted-F1 | ESMARG | 0.981 | 0.992 |
|  | DIAMOND | 0.732 | 0.991 |
|  | BLAST | 0.731 | 0.959 |
|  | ARG-SHINE | 0.806 | - |

“-” indicates that the model does not have this function.

### 3.3 Diversity of the resistomes in Lake Taihu

To determine the resistomes of CA, we applied ESMARG on the metagenome sequences of 26 CA samples from the northern bays of Lake Taihu where cyanobacteria carried out a near full-year bloom cycle. In our previous studies, a total of 610 Mb of 16S rRNA gene amplicon sequence data were generated from Illumina MiSeq sequencing, and 199 Gb of high-quality MG sequence data were generated from Illumina HiSeq sequencing [28, 29]. A total of 110 different ARGs, relevant to 24 drug classes, were identified (Table S1). To assess seasonal distribution, ARG abundance was normalized to the ARG copy number per bacterial cell [38]. The three most abundant ARGs were bacA (9.6%), macA (8.3%) and vanY (6.2%), which respectively confer Bacitracin, MLS, and Vancomycin resistance (Fig. 3A). The recently emerged ARG optrA [39] was also detected in CA samples at a percentage of 0.24%.

**Figure 3.**
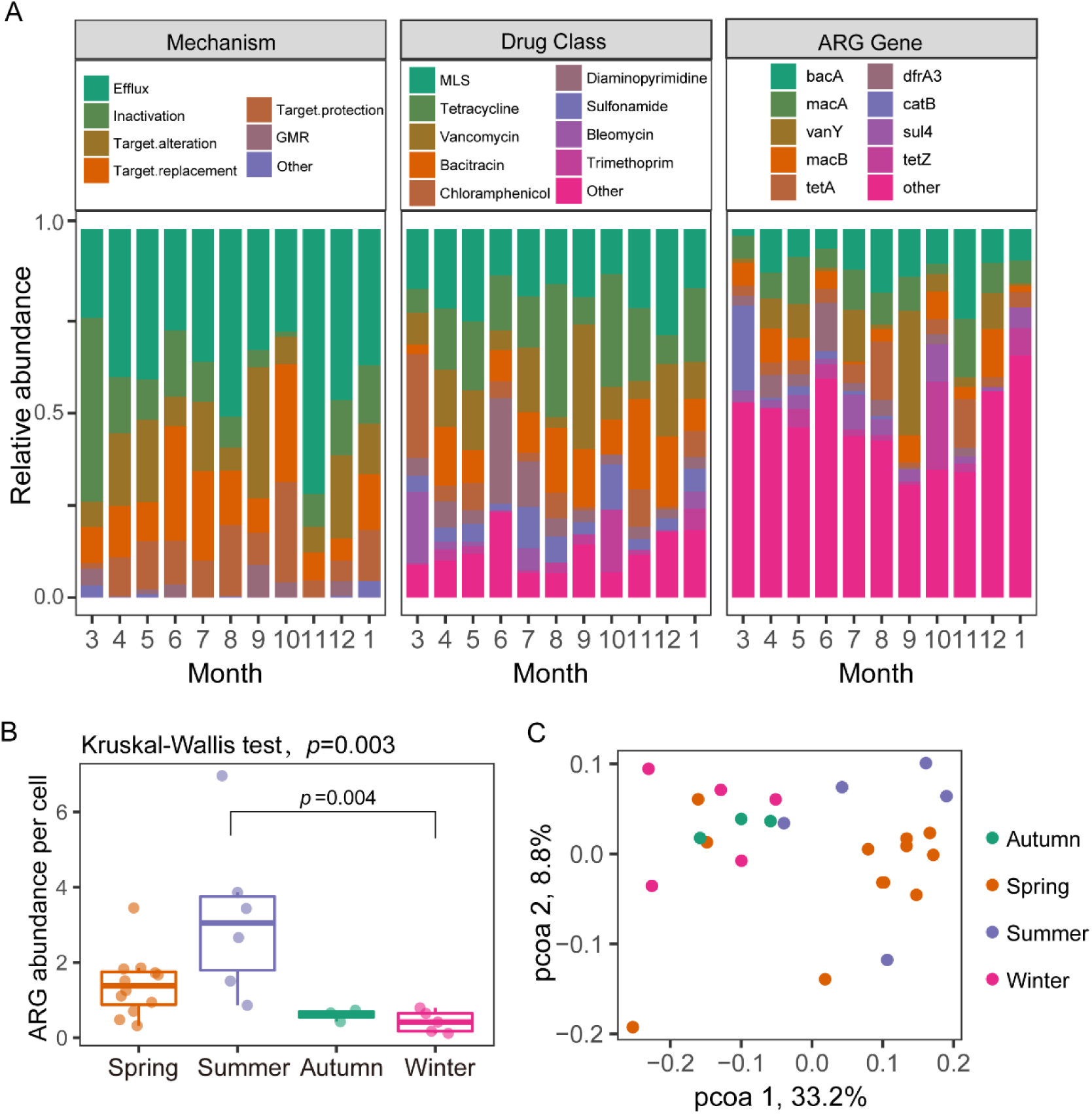
The abundance and seasonal variation of CA resistomes. **A)** Average relative ARG abundance across different sampling months based on resistance mechanism, drug class, and the nine most abundant ARGs. GMR: gene modulating resistance. MLS: Macrolides-lincosamides-streptogramins. **B)** Boxplots of the total ARG abundance (copy of ARGs per cell). Richness of the ARGs. Richness index was calculated based on a rarified matrix of resistance gene counts, which was subsampled to the lowest sample’s level. In the boxplots, hinges show the 25, 50, and 75 percentiles. The upper whisker extends to the largest value no further than 1.5 * IQR from the upper hinge, where IQR is the inter-quartile range between the 25% and 75% quartiles; The lower whisker extends to the smallest value at most 1.5 * IQR from the lower hinge. **C)** Principal coordinate analysis (PCoA) reveals distinct ARG composition diversity in four seasons.

Since different ARGs might be associated with the same resistance mechanism or drug class, the relative abundances of ARGs were aggregated based on their resistance mechanisms and drug classes. ARGs encoding antibiotic efflux were the most abundant, accounting for about 38.2% of the total ARG abundance. The next most prevalent were ARGs for and antibiotic inactivation (17.1%) (Fig. 3A). When ARGs were aggregated by drug class, ARGs conferring resistance to MLS (19.2%), Tetracycline (17.0%), and Vancomycin (12.5%) were the most abundant (Fig. 3A). The relative abundances of ARGs encoding major resistance mechanisms or drug classes varied significantly across samples.

### 3.4 Seasonal variation of the resistomes in Lake Taihu

The total ARG abundance showed significant difference across four seasons (Fig. 3B; *p* = 0.003, Kruskal-Wallis test). The mean ARG abundance was the highest in summer (3.21±2.16 copies per cell) and lowest in winter (0.43±0.29), and their means were significantly different (*p.adj* = 0.004, Dunn *post hoc* test). However, the mean ARG richness (Fig. S2A) and Shannon’s H index (Fig. S2B) showed no significant difference across four seasons. ARG abundance also varied across different months (*p* = 0.02, Kruskal-Wallis test): Jan. (0.39 ± 0.27), Oct. (0.42 ± 0.00), and Dec. (0.48 ± 0.44) were the lowest in mean ARG abundance, while Jun. (4.75 ± 1.92), Aug. (1.76± 0.86), and Apr. (1.72±0.90) were the highest (Fig. S2C). However, *post hoc* analysis indicated that total ARG abundance was not significantly different between any month pairs (*p.adj > 0.05,* Dunn *post hoc* test).

To identify structural differences of CA resistomes across seasons, PERMANOVA (Permutational multivariate analysis of variance) was performed at the individual gene level (Table 3). The resistomes exhibited significant differences (*p* < 0.05) when any two seasons were compared, with the exceptions of the comparisons between spring and summer, as well as autumn and winter. Clustering analysis at ARG gene level and drug-class level both showed a strong separation, with almost all samples of spring and summer clustered together, whereas samples from autumn and winter formed another cluster (Fig. 3C and Fig. S2D, E).

**Table 3.** Differences of the CA resistomes between seasons. PERMANOVA (Permutational multivariate analysis of variance using distance matrices) was performed based on Bray-Curti’s dissimilarity matrix at the level of individual ARGs. The upper triangle (shaded grey) shows the F values of PERMANOVA, while the lower triangle shows the *p* values with those less than 0.05 bolded.

|  | Spring | Summer | Autumn | Winter |
| --- | --- | --- | --- | --- |
| Spring |  | 1.28 | 2.04 | 3.27 |
| Summer | 0.207 |  | 2.28 | 4.13 |
| Autumn | 0.004 | 0.033 |  | 1.06 |
| Winter | 0.008 | 0.006 | 0.452 |  |

### 3.5 Relationship between resistomes and microbiome

To understand the relationships between resistomes and bacterial community structure, we performed Procrustes analyses (Fig. 4A). Procrustes analysis yielded a matrix-matrix correlation coefficient of 0.74 for metagenome 16S-based bacterial community structure (protest, *p* < 0.001), suggesting a strong association between CA bacterial community structure and the resistomes. Focusing on the relationship between resistomes and cyanobacteria, we also performed Procrustes analyses based on the cyanobacterial community structure obtained from semi-quantitative *in vivo* monitoring. The analysis also revealed a strong association with a matrix-matrix correlation coefficient of 0.61 and *p* = 0.003 (Fig. 4B).

**Figure 4.**
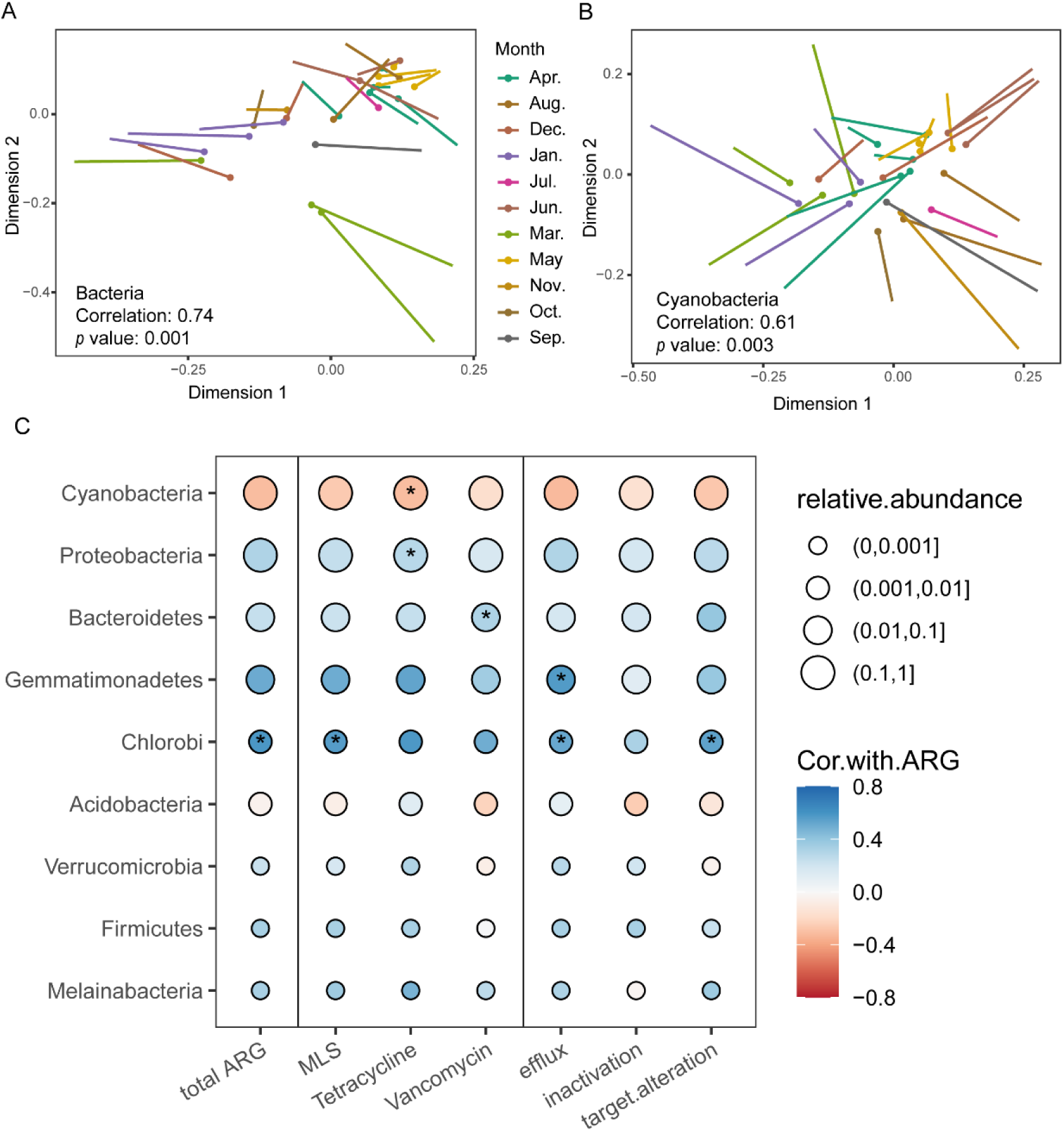
The linkage of the CA resistomes to microbiomes. **A**) Relationships detected by Procrustes analysis between the resistomes and bacterial community structure. Vegan Procrustes test ‘protest’ yielded a matrix-matrix correlation coefficient of 0.74 (protest, p < 0.001). **B**) Relationships detected by Procrustes analysis between the resistomes and cyanobacterial community. Vegan Procrustes test ‘protest’ yielded a matrix-matrix correlation coefficient of 0.61 (protest, p = 0.003). The dotted ends of lines represent the resistome position, while the undotted ends represent the bacteriome position. **C**) The association between the ARG abundance (total ARG abundance, top three major drug class and resistance mechanism) and the relative abundance of major bacterial phyla. The circle-filled color corresponds to Spearman’s correlation coefficient. The asterisks ‘*’ denote significant correlations (*p* < 0.05 after adjustment for multiple testing).

To further determine whether the relationships between the resistomes and microbiomes depend on phylogenetic lineages, we determined the linkages of the total ARG abundance, the top three major ARG drug classes and resistant mechanisms to the relative abundances of major phyla (Fig. 4C). The ARG abundances were positively correlated with Chlorobi (Spearman’s rho = 0.58, adjust *p* =0.02), and not positively correlated to any phyla. As exemplified by the abundances of MLS-, Tetracycline-, and Vancomycin-resistant genes, the top three major ARG classes, they were correlated with different phyla. Cyanobacteria, the most abundant phylum, were negatively correlated with ARG abundances related to Tetracycline (rho = -0.31, adjust *p* =0.001). In contrast, Proteobacteria, the second most abundant taxon, was positively correlated with ARG abundances related to Tetracycline (rho = 0.28, adjust p =0.001). In addition, since the total ARG abundance was highly correlated with ARGs of different resistance mechanisms (Pearson’s r = 0.35-0.91), or most classes (r = 0.40-0.88) (Fig. S3), results of the major ARG groups were largely similar to those based on the total ARG abundance. These results suggest that the resistomes in AS could be strongly tied to microbial physiology.

### 3.6 Biotic and abiotic drivers of CA resistomes

We performed a range of additional statistical analyses to help unravel the drivers of CA resistomes. First, we examined the linkage between resistomes and individual environmental variables, and KEGG pathway at level 1 (Table 4). Of the 15 environmental variables examined, seven of them had significant correlations with changes of ARG abundance (Table 4). Biochemical oxygen demand (BOD) was the best predictor of the ARG abundance (R= -0.67, *p* = 0.0001; Fig. 5A), as ARG abundance decreased with BOD. The dissolved oxygen (DO) also showed a negative correlation with the ATG abundance (R= -0.57, *p* = 0.001; Fig. 5B). In contrast, the temperature of water (WT) was positively correlated with the ARG abundance (R = 0.50, *p* = 0.005; Fig. 5C). Five functional gene classes at KEGG pathway level 2 were detected to significantly correlated with changes of ARG abundance (Table 4). Interestingly, ARG abundances in CA had strong positive relationships with genes for replication and repair (R=0.65, *p* = 0.0002; Fig. 5D) and nucleotide metabolism (R=0.40, *p* = 0.02; Fig. 5E), and the correlation was most negative with genes for Membrane transport (R=-0.52, *p* = 0.007; Fig. 5F).

**Figure 5.**
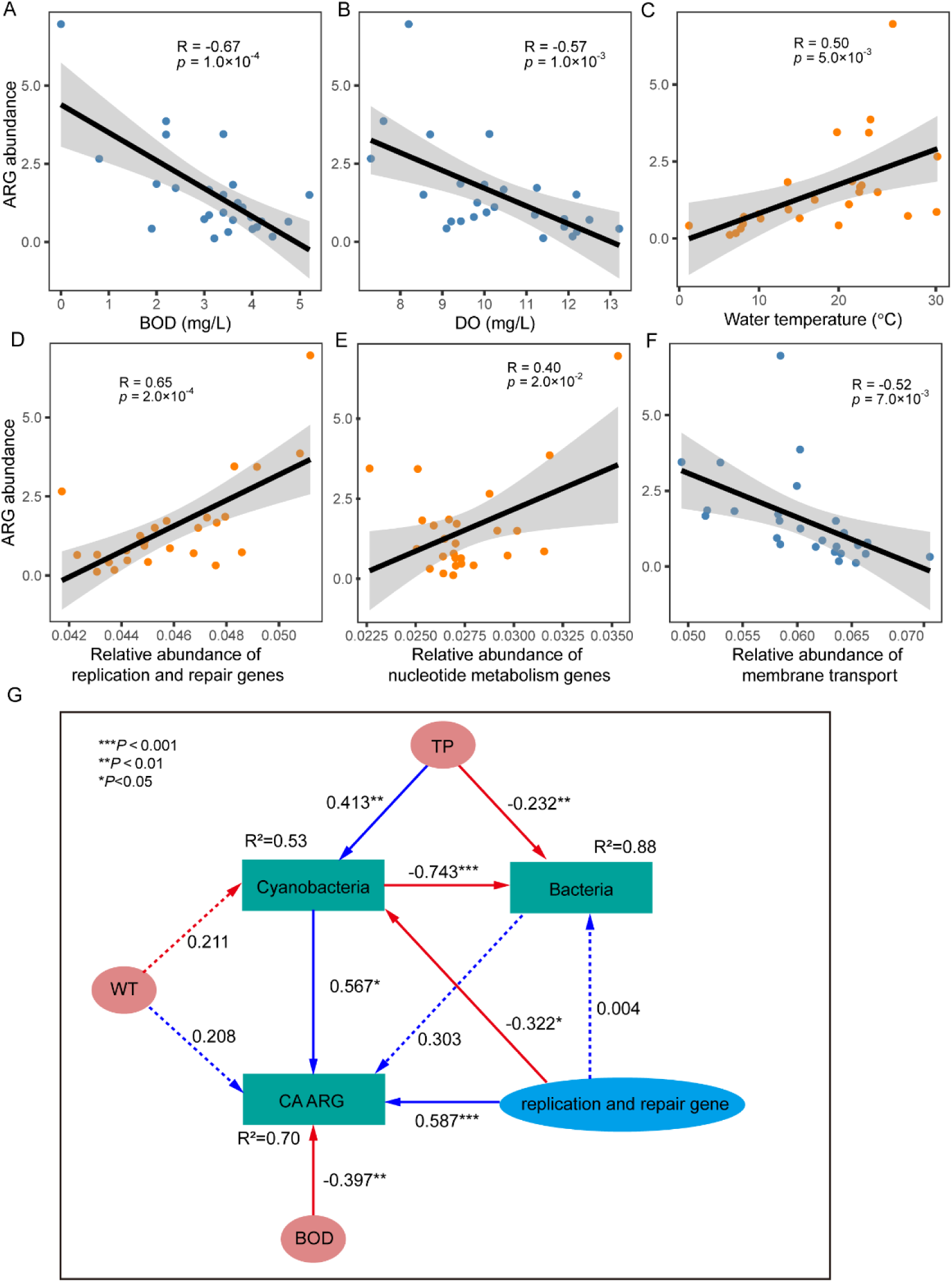
Drivers for the CA resistomes. **A-C)** Relationships of the total ARG abundance and environmental variables. **D-F)** Relationships of the total ARG abundance and functional variables. **G)** A structural equation model showing the direct and indirect effects of multiple environmental and functional variables on the total ARG abundance, cyanobacterial and bacterial diversity in CA. Environmental and functional variables are represented by circle. Composite variable represented by rectangles. Blue and red arrows represent significant positive and negative pathways, respectively. Numbers near the pathway arrow indicate the standard path coefficients. R^2^ represents the proportion of variance explained for every dependent variable. The goodness of fit was acceptable: Model χ^2^ = 5.79, df = 3, *p* = 0.122.

**Table 4.**
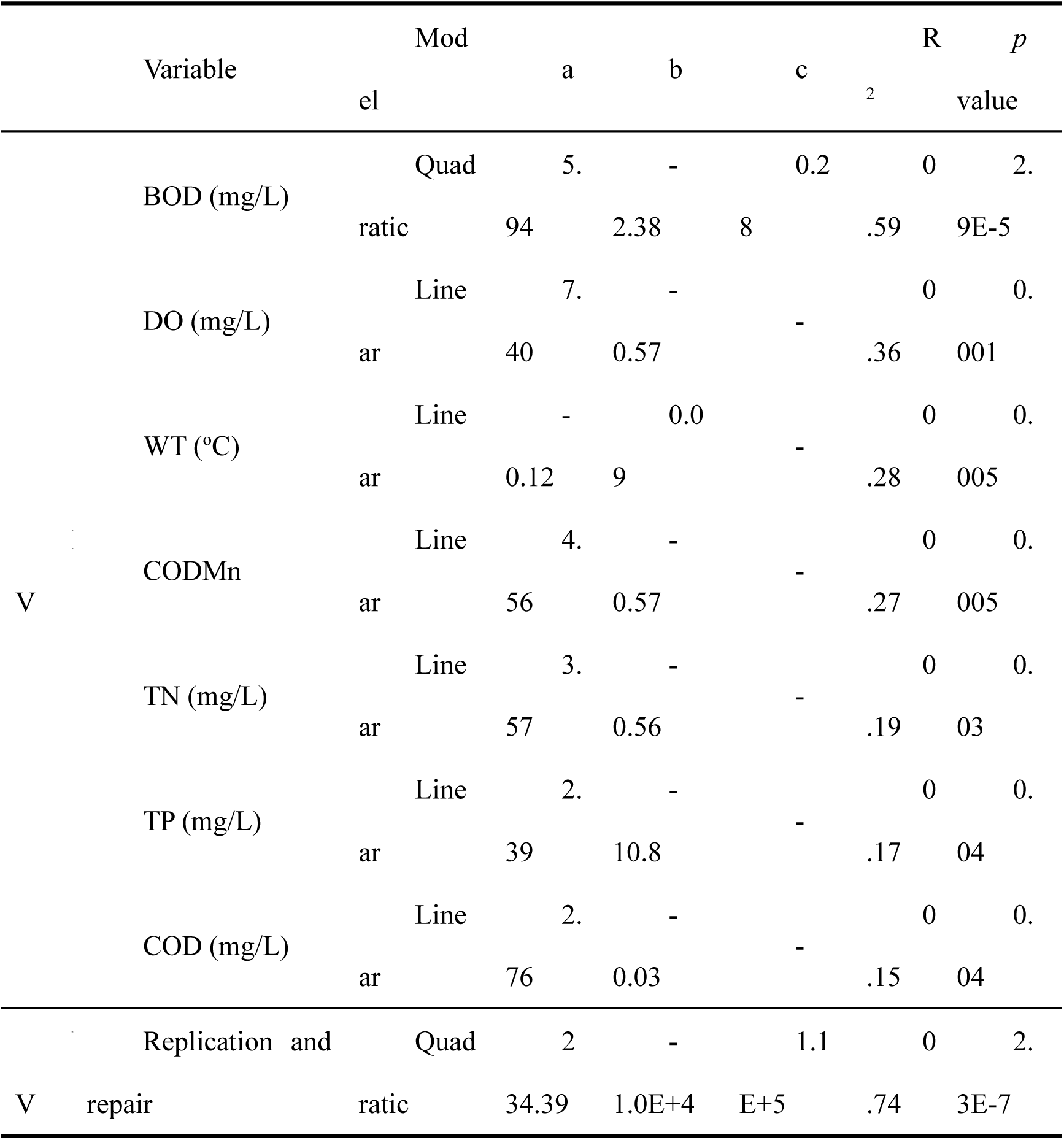
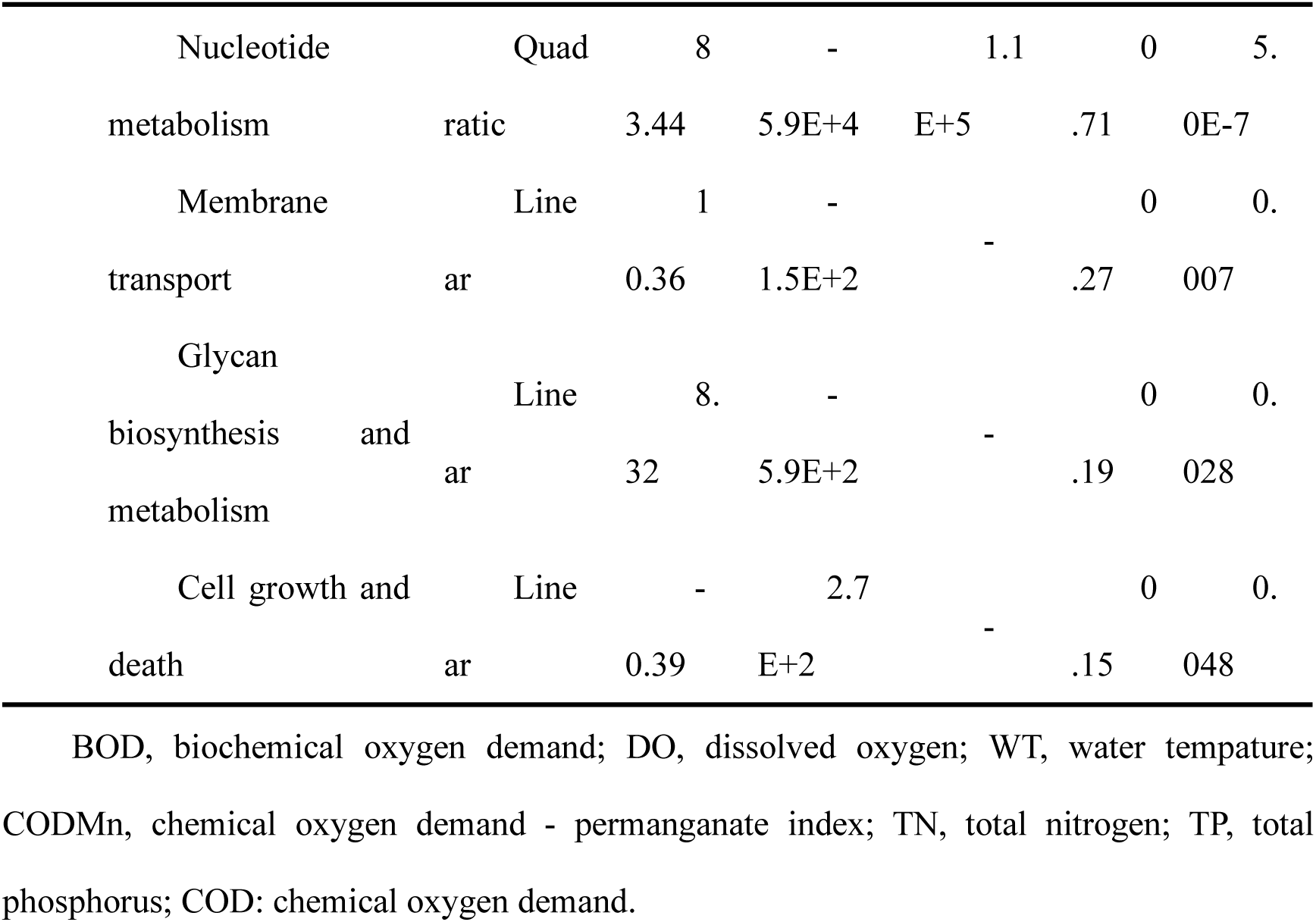
Univariate models predicting the total ARG abundance (ARG copies per cell) as a function of various environmental and functional variables. For each variable, we fitted a linear (y = a + bx) and a quadratic (y = a + bx + cx^2^) model and results are shown for the model with lower Akaike information criteria (AIC) value.

Structural equation model (SEM) analysis was further performed to understand the detailed relationships among ARG (copy number per bacterial cell), cyanobacteria, other bacteria and the correlated environmental and functional variables (Fig. 5G). The results show that the CA resistome was more correlated to cyanobacteria (standardized path coefficient, *β* = 0.57, *p =* 0.041) and bacterial functional variables (replication and repair: *β* = 0.58, *p* < 0.001) than environmental factors (BOD: *β* = -0.40, *p* = 0.002, WT: *β* = 0.21, *p* = 0.143). As expected, the cyanobacterial community was found to be the most influential variable directly related to bacterial diversity (*β* =-0.743, *p* < 0.001). The SEM result was consistent with the Procrustes analysis. These results suggest that the CA resistome were closely correlated with cyanobacterial community. Additionally, the attached bacteria in CA were more affected by the community composition of cyanobacteria than by environmental factors. Finally, the SEM model explained a high proportion (70%) of the resistome variations, which is mainly due to including the predictor of cyanobacteria.

## 4. Discussion

### 4.1 Advantages of ESMARG in ARG identification and classification

Based on our simulated test set and the Oxford dataset, ESMARG outperforms traditional sequence alignment tools like BLAST [11] and DIAMOND [12] and deep learning model ARGNet[18], demonstrating stronger generalization capabilities with higher precision and recall (Fig. 2B). As shown in Fig. 2A and Table 1, expanding the negative sample set improved the performance of ESMARG-1 in ARG identification. To further evaluate the impact of negative sample selection, we varied the DIAMOND threshold (from 1e-1 to 1e-10) to reconstruct negative sample sets and retrain the models (Fig. S1A). Fig. S1B shows that different filtering thresholds had limited influence on model performance (F1: 0.956–0.968), with the optimal performance achieved at a threshold of 1e-5.

ESMARG-1 demonstrates the capability to identify non-homologous ARGs. On the subset of 157 sequences in the test set that could not be aligned to the training set via DIAMOND (serving as non-homologous sequences), ESMARG-1 achieved an accuracy of 52.2% (82/157), outperforming ARGNet’s 49.7%, which highlights ESMARG-1’s ability in recognizing non-homologous ARGs.

ESMARG-2 and ESMARG-3 also exhibit the ability to identify non-homologous sequences. In function and mechanism classification tasks, ESMARG successfully annotated all sequences, whereas BLAST only identified 1212/1218 and 1116/1212, and DIAMOND recognized 1804 and 1121 sequences, failing to classify non-homologous entries (Fig. S1C). Additionally, as shown in Fig. S1D, ESMARG demonstrates faster prediction speed.

### 4.2 Seasonal variations of CA resistomes and its role in CyanoHABs

Temperature emerged as a major driver of seasonal ARG dynamics (Fig. 5G). In the Yellow River, it explained 37% of ARG variation, with high-risk ARGs (e.g., tetM, mecA, and bacA) increasing with temperature [40]. Similar patterns were observed in peri-urban rivers [41].Elevated temperatures enhance microbial metabolism and ARG expression [42]., while low temperatures suppress transcription, as seen with reduced β-lactamase expression [43]. Temperature also modulates global transcriptional regulators (e.g., *marA*) [44] and heat shock proteins (e.g., *ibpB*) [45], potentially enhancing resistance [46], mutation rates and promote horizontal gene transfer [47].

Our SEM analysis showed that cyanobacteria significantly promoted ARG abundance in CA samples (Fig. 5G), with temperature also promoting cyanobacterial growth (β=0.211). Cyanobacterial blooms, common in summer, coincide with ARG peaks. This synchrony suggests cyanobacteria mediate temperature-driven ARG dynamics.Cyanobacteria may influence ARG levels by altering microbial communities and water chemistry (e.g.,DO, pH, light) [48], favoring attached bacteria adapted to bloom conditions [49]. Additionally, bloom conditions increase the abundance of potentially drug-resistant taxa (e.g., *Ascomycota*, *Thickettsia*) [50] and release cyanobacterial toxins (e.g., microcystin), which impose selective pressure on microbial communities and promote ARG dissemination [51].

### 4.3 The influence of BOD and DO on CA resistomes

Besides temperature, biochemical oxygen demand (BOD) and dissolved oxygen (DO) were also key environmental drivers of ARG abundance witg negative correlations (Fig. 5A, B). Low BOD suggests limited organic matter, which may reduce general microbial activity. However, ARG-harboring microorganisms may persist due to their metabolic flexibility and capacity to utilize alternative resources [52].

DO, a water quality indicator, supports aerobic microbies that can suppress ARG carriers through competition and predation [53]. Conversely, low DO favors anaerobic bacteria that are more likely to carry and transmit ARGs [54]. Adequate DO levels is therefore crucial for controlling ARG proliferation. The relationships between these environmental factors and ARG abundance reveals the influence of microbial community structure and function on antibiotic resistance in water bodies. By controlling these environmental factors, the abundance of ARGs can be reduced, thereby protecting the health of water body ecosystems.

### 4.4 The relationship of ARG and microbial functions

Analysis of microbial functional genes revealed strong correlations between ARG abundance and core metabolic pathways. Replication and repair genes, as well as nucleotide metabolism genes, were positively correlated with ARG levels, while membrane transport genes were negatively correlated (Fig. D-F). Replication and repair genes are involved in the processes of DNA replication and repair. During DNA replication, the movement of the replication fork may be affected by environmental stresses such as antibiotics, leading to DNA damage, thereby introducing new mutations and generating new ARGs during the repair process. [55]. Homologous recombination may also facilitate the spread of ARGs among microorganisms [56]. Nucleotide metabolism genes, which mediate the synthesis and breakdown of nucleotides, are also closely linked to ARG expression. ARGs share intermediates or enzymes with these pathways, [57] and their upregulation under antibiotics can further promote ARG expression [58].

In contrast, membrane transport genes enable the influx and efflux of compounds across cell membranes. An increase in these genes may indicate enhanced efflux of antibiotics, reducing intracellular selective pressure and the need for ARG acquisition [59].Thus, microbial responses to stress may promote or reduce ARG abundance based on gene expression patterns. These findings emphasize the importance of integrating microbial functional gene analysis into ARG risk assessments and environmental monitoring efforts.

### 4.5 The influence of TN and TP on CA resistomes

Total nitrogen (TN) and total phosphorus (TP) are important nutrients and environmental driving factors of ARGs (Table 4 and Fig. 5G). Elevated TN and TP levels lead to eutrophication and intensified microbial competition and interactions., which can facilitate ARG enrichment and horizontal gene transfer within microbial communities [60]. Excessive nutrient inputs also lead to CyanoHAB [61], which can enhance ARG dissemination through community shifts and toxin production[62, 63]. Therefore, controlling TN and TP discharge is critical for preventing eutrophication and mitigating ARG-related environmental risks.

## 5. Conclusions

Collectively, our findings highlight the advantages of the ESMARG framework for ARG detection and classification in complex environmental samples. Seasonal patterns in ARG abundance are closely linked to temperature, microbial functional responses, and cyanobacterial activity. Key environmental drivers—including BOD, DO, TN, and TP—further shape the resistome by influencing microbial community structure and metabolic activity. These results underscore the importance of incorporating environmental context and microbial function into ARG monitoring strategies, and offer valuable insights for mitigating the ecological and public health risks associated with antibiotic resistance in freshwater ecosystems.

## Availability of data and materials

The DNA sequences of the metagenomes are deposited in the National Center for Biotechnology Information (NCBI) under the project accession number PRJNA664299.

The Python script for ESMARG is publicly available on GitHub in the YangLab-BUPT/ESMARG repository https://github.com/YangLab-BUPT/ESMARG.

## Authors’ contributions

HL and MQ drafted the work and contributed to the creation of new software used in the work.

XW, HL and QY contributed to the interpretation of data. TC and RJ contributed to the acquisition.

CZ and YY substantively revised the draft.

All authors have approved the submitted version and have agreed both to be personally accountable for the author’s own contributions. All authors read and approved the final manuscript.

## Ethics approval and consent to participate

Not applicable

## Consent for publication

Not applicable

## Supplementary Figures and Tables

**Figure S1.**
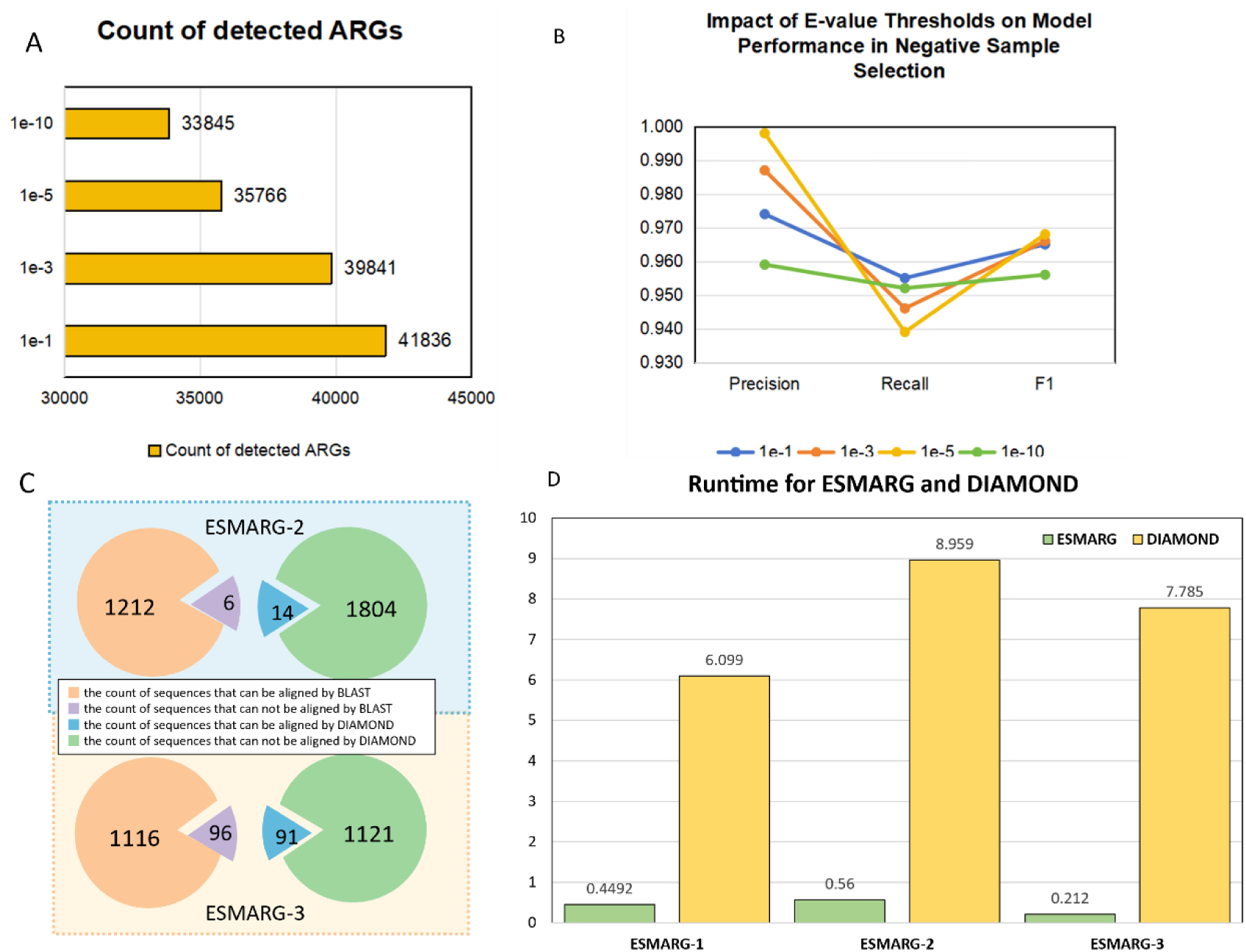
Utilized datasets and performance of ESMARG model. **A)** Negative sample screening using BLAST with different e-value thresholds. **B)** Model performance with negative samples selected by different e-value thresholds. **C)** BLAST and DIAMOND are limited by the aligned non-homologous sequence. **D)** Time comparison for DIAMOND with ESMARG-1, ESMARG-2 and ESMARG-3.

**Figure S2.**
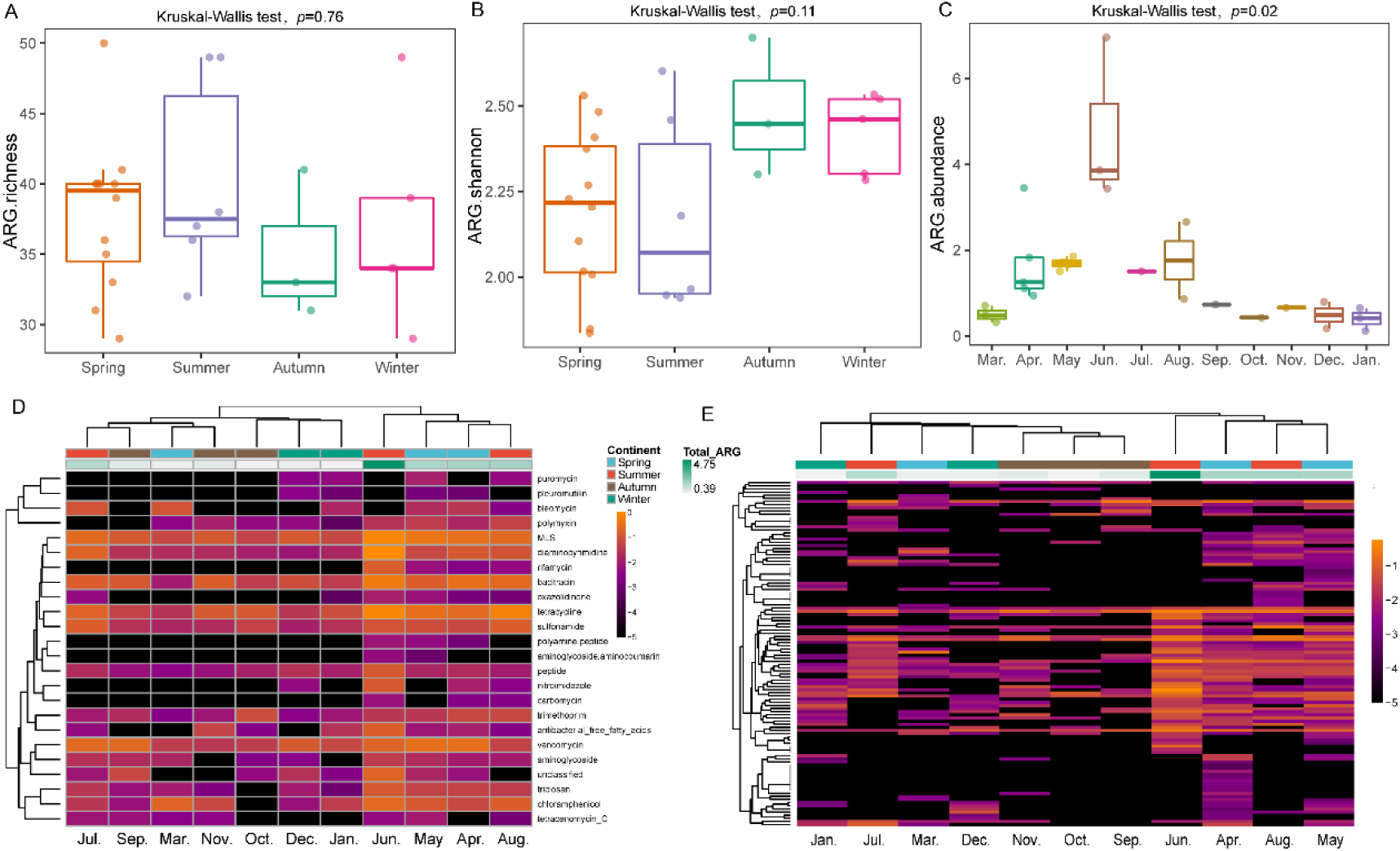
The composition and alpha diversity of ARGs across global CA samples. **A)** Boxplots of the ARG richness across four seasons. **B)** Boxplots of the ARG Shannon index across four seasons. Richness and Shannon index were calculated based on a rarified matrix of resistance gene counts, which was subsampled to the lowest sample’s level. **C)** Boxplots of the ARG abundance (copy of ARGs per cell). In the boxplots, hinges show the 25, 50, and 75 percentiles. The upper whisker extends to the largest value no further than 1.5 * IQR from the upper hinge, where IQR is the inter-quartile range between the 25% and 75% quartiles; The lower whisker extends to the smallest value at most 1.5 * IQR from the lower hinge. **D)** Cluster heatmap for the CA resistomes across months at drug class level. **E)** Cluster heatmap for the CA resistomes across months at gene level. Each column represents a month, which was calculated as the mean across samples within the month. Colors represent log-transformed gene abundance (copies of ARGs per cell). Average-linkage clustering of Manhattan distances was used to hierarchically cluster both months and drug classes or genes.

**Figure S3.**
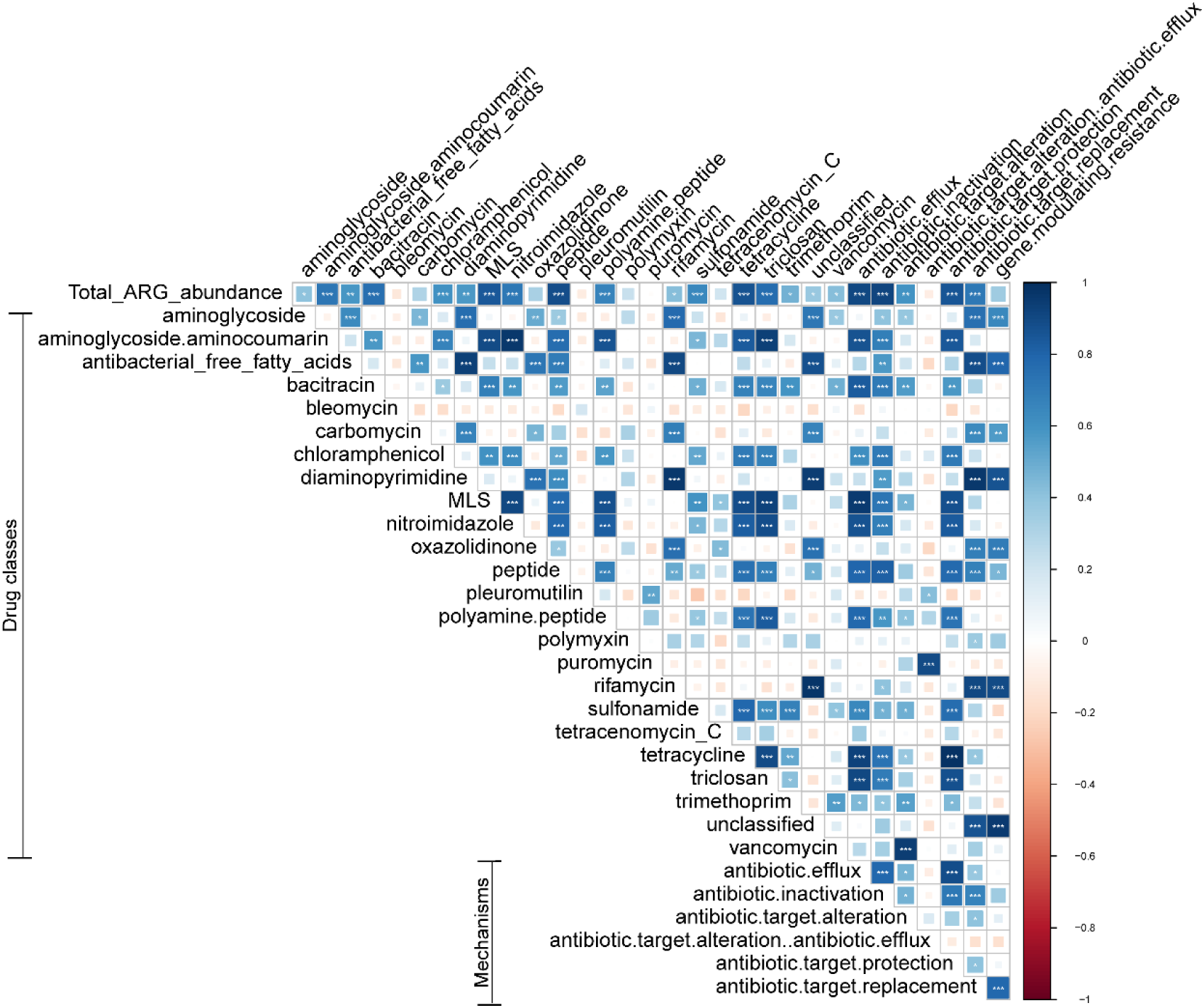
Pairwise correlations with respect to the total ARG abundance, drug classes, and resistance mechanisms. The color gradient indicates Pearson’s correlation coefficients, with more positive values (dark blue) indicating stronger positive correlations. The asterisks ‘*’ denote the significance levels of the Pearson’s correlation coefficients (n = 26 biologically independent activated sludge samples): *** p < 0.001, ** p < 0.01, and * p < 0.05 after adjustment for multiple testing.

