## Supplementary material for "Large-language-model-based antibiotic resistance gene prediction and resistomes mining in cyanobacterial blooms": appendix.pdf

#### Detailed dataset preprocessing

For ESMARG-1, a total of 8,733 ARGs were selected from MEGARes[1], 3,204 from ResFinder[2], and 12,746 from SARG[3]. After removing duplicates, the final ARG database included 15,624 unique ARGs. Negative samples for ESMARG-1 were selected from two primary datasets: the UniProt[4] database and GMGC[5]. For the UniProt database, only reviewed proteins, those with experimentally validated functions were included. Next sequences associated with ARGs were removed from the downloaded UniProt data, resulting in a set of 548,151 reviewed proteins. Additionally, the negative samples, non-ARG sequences, were collected from the GMGC database including the real environmental metagenomic sequences of food, livestock, compost, soil, and water. BLAST was employed to align GMGC sequences against known ARGs, using an e-value threshold of 1e-5. Then sequences identified as potential ARGs were removed and the remaining sequences formed the negative samples including 79,955 sequences. For testing, a set of 20,009 negative samples was chosen from UniProt and 3219 positive samples were selected from ARG database, ensuring no overlap with the training samples.

For ESMARG-2, the same training data as the DeepARG was utilized[6], encompassing 40 distinct function categories. The train and test sets were split with a ratio 8:2 for each category. For category with fewer than six sequences, only one sample was allocated to the test set. Finally, 9786 ARGs from 40 categories were involved in the training process, 2470 ARGs from 30 categories were included in test dataset, since 10 categories of ARGs had only one samples, the top 10 count of categories are shown in Fig. S1A. For ESMARG-3, the module designed for antibiotics mechanisms prediction, 6 categories of antibiotics mechanisms with 5422 sequences were collected in the training set and the test datasets contained 606 sequences.

#### Model hyperparameters

Modules ESMARG-1, ESMARG-2 and ESMARG-3 all utilized ESM1v to extract semantic features from sequences, obtaining the mean representations from the 33rd layer of ESM1v[7] for further training (shown in Fig. S1). ESMARG-1 comprised three layers, interconnected by dropout layers (dropout rate = 0.5). The first layer transformed the input mean representation with 1280 dimensions into a semantic space with 256 dimensions, followed by the application of a ReLU activation function. The second layer includes 64 units. The third layer condensed the output to a single value, utilizing a Sigmoid activation function to generate probability score for binary classification. The learning rate for ESMARG-1 was set to 5e-4.

ESMARG-2 module comprised an input layer with 1280 units and two hidden layers consisted of 128 and 64 units, respectively. A ReLU activation function was applied to generate the final output. The output layer of ESMARG-2 consisted of 40 units, corresponding to the 40 function categories of ARGs. ESMARG-3 was constructed with an input layer of 1280 units, followed by two hidden layers with 256 and 64 units. A ReLU activation function was employed to generate the final output. The output layer of ESMARG-3 consisted of six units, corresponding to six categories of antibiotic resistance mechanisms. The loss function of ESMARG-3 was the Focal Loss, which addressed the imbalance in the number of resistance mechanism categories. ESMARG-4 employed DIAMOND directly to perform sequence alignment among ARG sequences predicted from the ESMARG-1 module, with parameters set to an e-value of 1e-5 and a minimum identity threshold of 20%.

#### Statistical analyses

Kruskal–Wallis and the Dunn *post hoc* tests were used to compare the means of ARG abundance across four seasons and their differences, using R function 'kruskal.test' and function 'dunnTest' of *FSA* R package [8]. Richness and Shannon's H index were computed using the *vegan* R package [9] to measure the diversity of ARGs based on a rarified count matrix, which was obtained by rounding the coverages and sub-sampling to the lowest sample's level. The heat map was generated to display the F1 score of different models on various ARG classifications using the function 'aheatmap' of *NMF* R package [10]. PERMANOVA was applied to identify structural differences of CA resistomes across seasons using the function 'adonis2' of *vegan* R package. Procrustes analysis was performed to test the relationships between resistomes and bacterial community structure using the function 'procrustes' of *vegan* R package in which the ordinations of the bacterial taxonomic composition and the resistome were generated from PCoA.

Structural equation modeling (SEM) [11] was used to explore the detailed relationships between multiple environmental and functional variables with the ARG (copy number per bacterial cell), cyanobacterial, and bacterial diversity in CA. We initially considered a full model that included all possible pathways (involving 7 environmental variables and 5 functional variables), then sequentially eliminated non-significant pathways until we generated a final model where all pathways were significant, leaving behind environmental variables TP, WT, BOD, and functional variables for replication and repair genes. We used the chi-squared test ($\chi^2$ test) [12] and root mean square error of approximation (RMSEA) and calculated the coefficient of determination $R^2$ [13] to assess how well the model fits . The SEM-related analysis was performed using the R package 'lavaan' [14].

### Figure

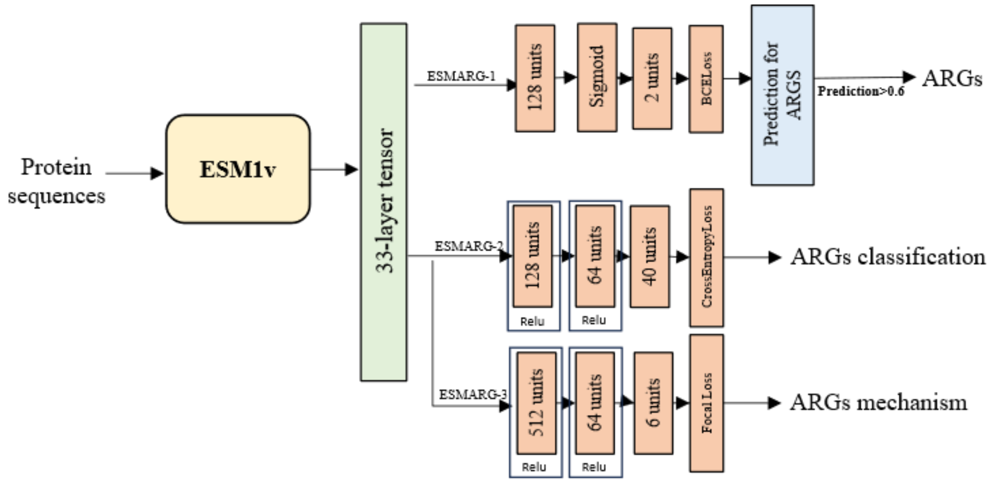


#### Figure S1. ESMARG's model structure

Detailed Model structure of ESMARG. ESMARG-1, ESMARG-2 and ESMARG-3 all use ESM1v's 33-layer tensor as feature and use MLP to train the training set.
